# Extracellular histones stimulate collagen expression and potentially promote liver fibrogenesis via TLR4-MyD88 signalling pathway

**DOI:** 10.1101/2020.09.17.302240

**Authors:** Zhi Wang, Zhenxing Cheng, Simon T Abrams, Ziqi Lin, ED Yates, Qian Yu, Weiping Yu, Pingsheng Chen, Cheng-Hock Toh, Guozheng Wang

## Abstract

**BACKGROUND:** Liver fibrosis progressing to liver cirrhosis and hepatic carcinoma is very common and causes more than one million deaths annually. Fibrosis develops from recurrent liver injury but the molecular mechanisms are not fully understood. Recently, the Toll-like receptor (TLR) 4-MyD88 signalling pathway has been reported to contribute to fibrosis. Extracellular histones are the ligands of TLR4 but their roles in liver fibrosis have not been investigated.

**AIM:** This study aims to investigate the roles and potential mechanisms of extracellular histones in liver fibrosis.

**METHODS:** *In vitro,* the LX2 cells, a human hepatic stellate cell (HSC) line, were treated with histones in the presence or absence of non-anticoagulant heparin (NAHP) for neutralising histones or TLR4-blocking antibody. The cells resultant expression of collagen I was detected using Western blotting and immunofluorescent staining. *In vivo*, the CCl_4_-induced liver fibrosis model was generated in male 6 week old ICR mice and in TLR4 or MyD88 knockout and parental mice. Circulating histones were detected and the effect of NAHP was evaluated.

**RESULTS:** Extracellular histones strongly stimulated LX2 cells to produce collagen I. The histone-enhanced collagen expression was significantly reduced by NAHP and TLR4 blocking antibody. In CCl_4_-treated wild type mice, circulating histones were dramatically increased and maintained high levels during the whole course of fibrosis-induction. Injection of NAHP not only reduced alanine aminotransferase (ALT) and liver injury scores, but also significantly reduced fibrogenesis. Since the TLR4-blocking antibody reduced histone-enhanced collagen I production in HSC, the CCl_4_ model with TLR-4 and MyD88 knockout mice was used to demonstrate the roles of the TLR4-MyD88 signalling pathway in CCl_4_-induced liver fibrosis. The levels of liver fibrosis were indeed significantly lower than in these knockout mice than the wild type parental mice.

**CONCLUSION:** This study demonstrated that extracellular histones are able to stimulate HSC to produce collagen I and TLR4 is involved in this process. The in vivo findings support the novel concept that high levels of circulating histones potentially stimulate TLR4 receptor to enhance fibrogenesis via the TLR4-MyD88 signalling pathway. NAHP detoxify extracellular histones and thus has a potential therapeutic role by reducing liver injury and fibrogenesis.

**Core tip:** This work fills the gap between recurrent liver injury and liver fibrosis. When liver cells die, histones will be released. High levels of extracellular histones not only cause a secondary liver injury, but also activate the TLR4-MyD88 signalling pathway to enhance collagen I production and liver fibrosis. Binding of non-anticoagulant heparin (NAHP) to extracellular histones reduces histone toxicity, alleviates liver injury and prevent histones from activating the TLR4-MyD88 signalling pathway. These results may explain why NAHP reduces liver fibrosis in this animal model although further investigations are required.

**ARTICLE HIGHLIGHTS:** *Research background:* Currently, the molecular mechanisms of liver fibrosis are not fully understood. Recurrent liver injury or inflammation initiate wound healing process along with fibrogenesis. However, what initiates this process is not clear. When cells die, damage-associated molecular patterns (DAMPs) will be released. Histones are the most abundant DAMP and are also the ligands for TLR4, which in turn has been demonstrated to be involved in bile duct ligation-induced liver fibrosis. Lipopolysaccharides (LPS) has been proposed to be a ligand for TLR4. Since recurrent liver injury does not naturally produce LPS but abundant extracellular histones, this study sought to investigate the potential roles of extracellular histones as TLR4 ligands in liver fibrosis.

*Research motivation:* Since our laboratory has been involved in studying the roles of DAMPs in critical illnesses, extracellular histones in liver fibrosis is of great interest in terms of biological and clinical significance.

*Research objectives:* Our study aimed to clarify the roles of extracellular histones in fibrogenesis in vitro and in vivo.

*Research methods:* In our study, a HSC cell line and animal models of liver fibrosis were employed. Intervention studies with NAHP to detoxify histones and TLR4 blocking antibodies to inhibit TLR4 was performed. In addition, TLR4 and MyD88 knockout mice were used to support that the theory that the TLR4-MyD88 signalling pathway is involved in liver fibrosis in the CCl_4_ mouse model.

*Research results:*

1. Demonstrated that high levels of circulating histones exist when fibrosis is induced by CCl_4_ in the mouse model.
2. Demonstrated *in vitro* that extracellular histones are able to stimulate HSC cells to produce collagen I.
3. Demonstrated that NAHP is able to inhibit histone-enhanced collagen production in the HSC cell line, and reduce liver injury and fibrosis in the animal model.
4. Demonstrated that TLR4 is involved in histone-enhanced collagen I production in HSC cells. *In vivo*, the TLR4-MyD88 signalling pathway was demonstrated to be involved in liver fibrosis, but whether circulating histones are the major activators of the pathway is not fully clear.

*Research conclusion:* Recurrent liver injury releases extracellular histones which potentially activate TLR4-MyD88 signalling pathway to promote liver fibrosis. The ability of NAHP to detoxify circulating histones holds the potential for the treatment of liver injury and prevention of liver fibrosis.

*Research perspectives:* Future studies demonstrating the contributions of circulating histones in activating the TLR4-MyD88 signalling pathway *in vivo* will validate their role in liver fibrosis. Development of better anti-histone reagents to reduce liver injury and prevent liver fibrosis will hold great potential in the management of diseases with recurrent liver injury.

## Introduction

Liver fibrosis progressing to liver cirrhosis and hepatic carcinoma (HCC) is very common, both in China due to high incidence of Hepatitis B and in Western countries due to alcohol consumption and Hepatitis C^[1–3]^. In 2017, liver cirrhosis caused more than 1.3 million deaths globally, and constituted 2.4% of total deaths^[4]^. However, elucidating the underlying molecular mechanism and the development of specific therapies for fibrosis and cirrhosis have progressed very slowly and significant discoveries in this area are urgently needed.

Liver fibrosis is a natural wound healing response to chronic liver injury, which has many common causes, including viral infection, alcohol abuse, autoimmune diseases, drug toxicity, schistosomiasis, and metabolic diseases^[5]^. This process can be easily reconstituted in animals by administering chemicals toxic to hepatic cells, including carbon tetrachloride (CCl_4_), thioacetamide, and dimethyl or diethyl nitrosamine^[6–8]^. Bile duct ligation (BDL) and animal models mimicking specific chronic liver diseases are also used ^[6]^. CCl_4_ is mostly commonly used to study the underlying molecular mechanisms and evaluate therapeutic reagents for inhibiting liver fibrosis in animals ^[6]^. CCl_4_ directly damages hepatocytes by ligation of CCl_3_ to plasma, lysosomal, and mitochondrial membranes, as well as to highly reactive free radical metabolites^[9]^. A single dose of CCl_4_ causes centrizonal necrosis and steatosis with recurrent administration leading to liver fibrosis ^[10]^.

The pathological feature of liver fibrosis is the extensive deposition of extracellular collagens produced by myofibroblasts^[1]^. In the healthy liver, no myofibroblasts are present but they accumulate in the damaged liver and play a pivotal role in fibrogenesis^[11]^. Myofibroblasts are potentially derived from different types of cells. Hepatic stellate cells (HSCs) in the peri-sinusoidal Disse space are recognised as the most important source^[12]^. Bone marrow also contributes to both macrophages and to HSCs within the injured liver^[13]^. Recurrent injury, chronic inflammation and other chronic stimuli can cause HSCs to evolve into a myofibroblast-like phenotype and produce an extracellular matrix^[14]^. The myofibroblast-like phenotype also develops and expresses α-SMA *in vitro* when quiescent HSCs were isolated and cultured over a period of 5-10 days^[15]^.

HSCs activated both *in vivo* and *in vitro* express Toll-like receptor-4 (TLR-4), CD14 and MD2, therefore lipopolysaccharides (LPS) are able to assemble a complex to activate TLR-4 on HSCs ^[16]^. TLR-4 signals via MyD88-dependent and TRIF-dependent pathways which can activate fibroblasts, to promote fibrogenesis in numerous organs, including the liver, lungs, kidneys, heart and skin^[17–20]^. Since circulating LPS is produced primarily from Gram negative bacterial infections^[21]^, ligands that activate the TLR-4 receptor in chronic liver injury without concomitant bacterial infection have not been identified as yet. Increased intestinal permeability allows LPS to enter the circulation in the later stages of liver cirrhosis, but this leakage has not been demonstrated in the early stages of fibrogenesis ^[22]^.

Liver cell injury and death also release damage-associated molecular patterns (DAMPs) which may initiate the wound-repair process^[23]^. Extracellular histones, the most abundant DAMP, have gained great attention in recent years, as they play key roles in many pathological processes and human diseases^[24–35]^. Histones are well-conserved proteins that are essential for DNA packaging and gene regulation^[36]^. During tissue damage and cell death, nuclear chromatin is cleaved into nucleosomes, which are released extracellularly ^[37]^ and further degraded into individual histones^[38]^. Although histones are rapidly cleared by the liver^[39]^ and rarely detected in blood^[25, 26, 31, 32]^, liver cell death is likely to release histones locally^[40–43]^, which stimulate adjacent cells, including hepatic stellate cells. Histones could activate TLR-2, TLR-4 and TLR-9 receptors^[24, 40, 43–46]^ in the early stage of chronic liver injury and fibrogenesis. Therefore, TLR-4 activation is more likely to be maintained by extracellular histones rather than LPS.

In this study, we investigate the roles and mechanisms of extracellular histones in liver fibrogenesis. Circulating histone levels were measured in a CCl_4_-induced mouse liver fibrosis model. Extracellular histones were used to stimulate human hepatic stellate cell lines to demonstrate its direct effect on collagen I production. In addition, we used non-anticoagulant heparin (NAHP) to neutralize circulating histones in both the cell culture system and mouse liver fibrosis model to explore the role of anti-histone therapy in reducing collagen I production and fibrosis. To clarify the potential molecular mechanisms, we used a TLR4 neutralizing antibody to block histone-enhanced collagen production by a hepatic stellate cell line and compared the fibrogenesis in TLR4 and MyD88 knockout mice with their wild type (wt) parental mice in response to CCl_4_ treatment.

## MATERIALS AND METHODS

### Cell culture

LX2 cells, a human HSC cell line, was purchased from Shanghai MEIXUAN Biological Sciences and Technology LTD and were routinely cultured according to the suppliers’ instruction.

### Histone treatment

After optimization of the doses and times, 0-20 μg/ml calf thymus histones (Sigma) were added to cell culture medium to treat the LX2 cell line. Fresh medium with different concentrations of histones were changed every 48h. At day 6, medium and cell lysates were collected for Western blotting.

### TLR-4 neutralising antibody and NAHP treatment

The neutralizing antibody, PAb-hTLR-2, was purchased from Invivogen USA. LX2 cells were treated with 5μg/ml histones and 5μg/ml antibody. NAHP was synthesised and characterised by Dr. Yates in the University of Liverpool. NAHP (25 and 50μg/ml) was used to treat LX2 cells together with 5μg/ml histones. The cell culture medium was changed every 48h. Then cell lysates of the treated LX2 were collected at day 6.

### Western blotting

One ml of culture medium was collected and proteins were precipitated using ice cold acetone and suspended in 50μl clear lysis buffer ((1% SDS, Glycerol, 120mM Tris-HCL, 25mM EDTA, H2O, pH 6.8). Cells were washed with phosphate-buffered saline (PBS) 3 times and lysed using clear lysis buffer. Protein concentrations were determined using Bio-Rad protein assay kit II. Both medium and cell lysates (20μg protein) were subjected to SDS-PAGE and electrically transferred onto Millipore PVDF membranes. After blocking with 5% milk in TBST buffer (50 mM Tris-Cl, pH 7.6; 150 mM NaCl, 0.1% (v/v) Tween-20), the membranes were incubated with the sheep anti-human collagen I alpha 1 antibody (1:2000, R&D systems) overnight at 4°C. After extensive washings with PBST, the membranes were incubated with rabbit anti-sheep IgG-HRP (1:10,000, Santa Cruz Biotechnology, Inc) for 1h at room temperature. The protein bands were visualised using ECL (Millipore). For beta-actin, primary antibody was purchased from Abcam and anti-rabbit IgG-HRP was purchased from Santa Cruz. The band density was analysed using Image J software and then the ratios of collagen I/ beta actin were calculated.

### Immunofluorescent Staining

LX2 cells were seeded in 8-well Chamber slides and treated with or without 5μg/ml histones for 6 days. The cells were fixed with 4% (w/v) paraformaldehyde for 10 min and permeabilized with PBS+1% (v/v) Triton X-100 for 10 min. After blocking with 5% (w/v) milk +2 % (w/v) BSA in PBS, the cells were incubated with sheep anti-human collagen I alpha 1 antibody (1:200) or rabbit-anti-α-SMA antibody (1:200, Proteintech, United States) at 4°C overnight. After extensive wash, the cells were incubated with anti-sheep-FITC (Abcam, 1:2000) or anti-rabbit FITC (Abcam, 1:2000) at room temperature for 45min. Following extensive washing, the slides were mounted on cover slips using mounting media and antifades (Thermo Fisher scientific, UK). The fluorescent images were recorded using LSM-10 confocal microscope (excitation: 488nm and emission: 512nm).

### Animals

Mice were housed and used in sterile conditions at the Research Centre of Genetically Modified Mice, Southeast University, Nanjing, China. The rats were kept in a temperature and humidity control laminar flow room with an artificial 10-14h light cycle. All mice had free access to tap water. Sample sizes were calculated using power calculation based on our previous data and experience. All procedures were performed according to State laws and monitored by local inspectors and also approved by the Animal Research Ethics Committee at the Medical School of the Southeast University. ZW and CZX hold the full animal license for use of mice (Jiangsu Province, 2151981, and 2131272).

### CCl_4_ model with ICR mice

Healthy male ICR mice, 6 weeks old, were purchased from Yangzhou University Animal Experiment Centre and divided randomly into 3 groups: (1) 5 μL/g of 25% (w/v) CCl4 in olive oil injected intraperitoneally (i.p), twice a week; (2) CCl_4_ +NAHP: on the day of CCl_4_ injection, 8 μl/g body weight NAHP (4μg/ml in saline, filtered for sterility) were injected subcutaneously (s.c) every 8h. On the remaining days, 8 μl/g NAHP were injected s.c every 12 hours. (3) Saline was injected i.p and NAHP injected s.c as control. Three mice in each group were monitored every day and euthanized by neck dislocation at 4h, 8h 24h after first the dose of CCl_4_ each week to collect the blood and organs for further analysis. No mice died before euthanization.

### CCl_4_ model with genetic modified mice

C57BL/6JGpt (Wild type), TLR-4-KO (B6/JGpt-Tlr4^em1Cd^/Gpt) and MyD88-KO(B6/JGpt-Myd88^em1Cd^/Gpt) mice were purchased from Gempharmatech (Nanjing, China). Each type of mouse was divided into 3 groups as above. Eight mice in each group were treated the same as above for 28 days. All mice were euthanized at day 28. Blood and organs were collected for further analysis. No mice died before euthanization.

### Tissue processing and staining

In our pathology laboratory, organ tissues were routinely processed and 4μm sections were prepared. Staining with hematoxylin and eosin (H&E) and Sirius red were routinely performed and α-smooth muscle actin (α-SMA) was detected by immunohistochemical staining using anti-α-SMA (Proteintech, United States) as described previously^[47]^. Liver histological grading according to a previous publication^[48]^ were performed by two investigators in a blinded manner. The staining areas of α-SMA and fibrosis were quantified using Image J software.

### Measurement of blood histones and ALT

Venous blood was collected with citrate anti-coagulant and immediately centrifuged at 1500xg for 25min at room temperature. Plasma was collected and alanine aminotransferase (ALT) levels were measured by an automatic blood biochemical analyser in the clinical laboratory and histone levels were measured by Western blotting using anti-histone H3 (Abcam) as described previously^[32]^. In brief, plasma and histone H3 protein (New England Biolabs) as standard were subjected to SDS-PAGE and electrically transferred onto PVDF membrane (Millipore). After blocking, the blot was probed by anti-histone H3 (Abcam, 1:2000) at 4°C overnight. After extensive washing, HRP-conjugated secondary antibody (Santa Cruz, 1:10,000) was incubated at room temperature for 1h. After further extensive washing, ECL (Millipore) was used to visualize the protein bands, which were quantified against histone H3 protein standard to obtain the concentration of histone H3 in plasma using Image J software. The total histones were estimated based on the molar ratio of H3 in the cell nuclei.

### Measurement of hydroxyproline in the liver tissues

Part of the liver from euthanized mice was washed with saline and frozen at −80°C. Two hundred and fifty milligrams of frozen liver tissues were ground using low temperature Tissuelyser (Shanghai Jingxin Industrial Development Co. Ltd) and hydroxyproline was measured using a colorimetric kit from Beijing Solarbio Science & Technology Co., Ltd according to the instructions from the supplier.

### Statistical analysis

Continuous variables are presented as mean ± standard deviation (SD). Differences in means between any two groups were compared using unpaired Student’s t test. Differences in means between more than two groups were compared using ANOVA test followed by Student-Newman-Keuls test. A P-value < 0.05 was considered significant. Statistical analysis was performed using SPSS (version 25).

## RESULTS

### High levels of circulating histones are detected in mice treated with CCl_4_

CCl_4_ is a hepatotoxin that directly damages hepatocytes. CCl_4_ administration in mice to generate liver fibrosis is the most commonly used animal model, but it is not clear whether high levels of histones actually exist under these conditions in the circulation. Using Western blotting, circulating histone H3 was detected and the total circulating histone levels were calculated based on these levels of H3 as described previously^[32]^. We found that histone H3 was detectable at 4 hours after CCl_4_ administration (Figure 1A). In mice treated with CCl_4_+non-anticoagulant heparin (NAHP), H3 was also detectable (Figure 1B). Total circulating histones in CCl_4_ group reached peak values around 8 hours, whilst CCl_4_+NAHP reached peak values around 24h after the first dose of CCl_4_ was administered. Circulating histones were then reduced levels prior to the subsequent dose of CCl_4_ (Figure 1C). Average levels of total histones during each week are shown in Figure 1D, these were comparable between the groups except at week 1. At this time, the CCl_4_ alone group had lower levels of histones than the group treated with CCl_4_+NAHP. In contrast, histone H3 was barely detectable in control group treated with saline +NAHP (Data not shown). To monitor liver injury, blood alanine aminotransferase (ALT) was measured using a clinical biochemistry setting. The means±SE from 9 mice at day 28 were compared and it was found that NAHP alone did not increase ALT levels, whilst CCl_4_ caused a significant elevation of ALT (about 10 fold), but this elevation was significantly reduced by NAHP (about 40% reduction) (Figure 1E). The liver injury scores of haemotoxylin and eosin (H&E)-stained liver sections were also significantly reduced by NAHP treatment. These observations indicated that NAHP treatment did reduce liver injury.

**Figure 1.**
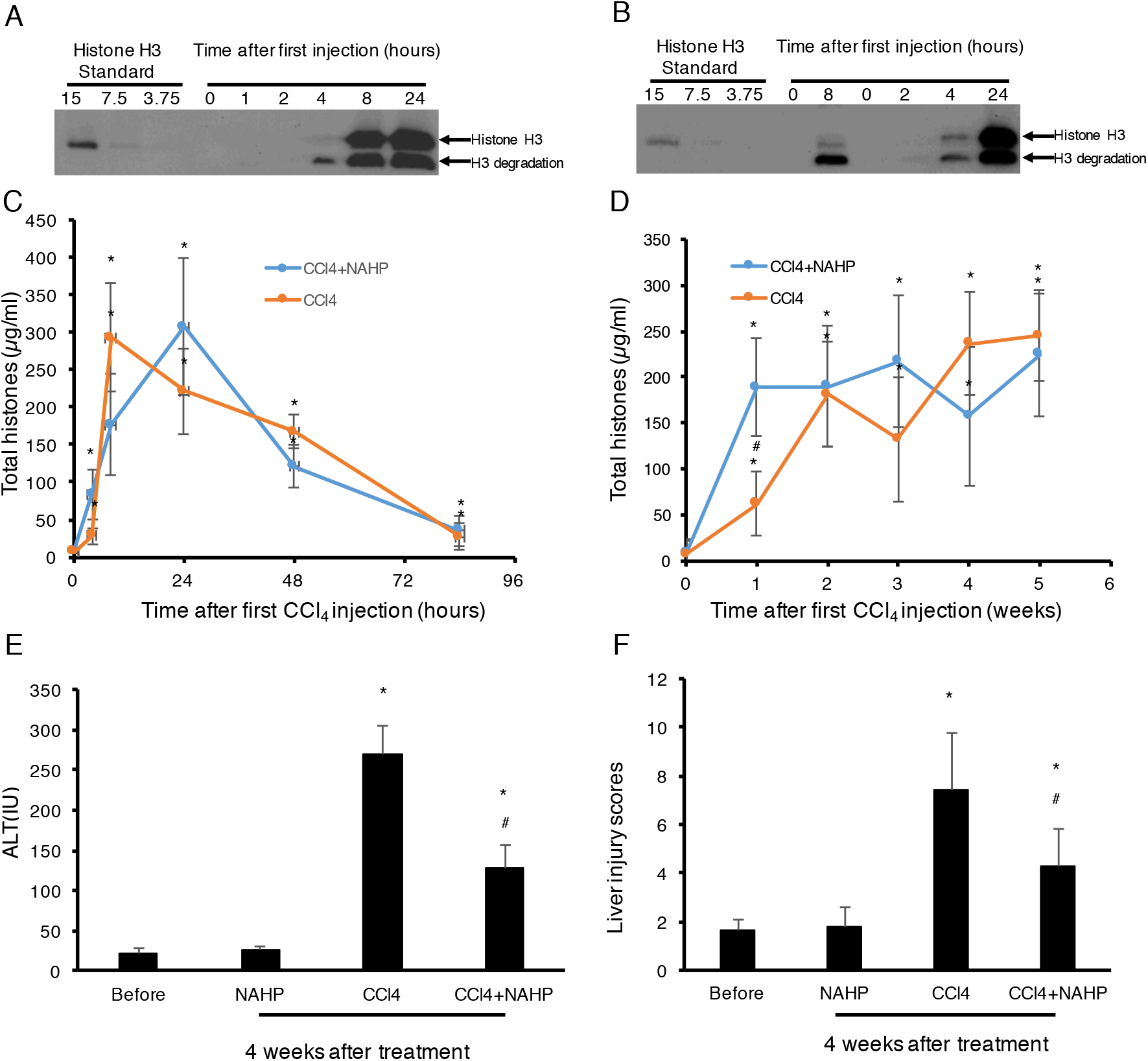
Circulating histones elevated in mice treated with CCl_4_. Typical Western blots of histone H3 standard (μg/ml) and histone H3 in plasma from mice treated with the first dose of CCl_4_ are presented (A) and CCl_4_+NAHP (B). The means±SD of circulating histones at different time points after first dose of CCl_4_ (Orange) and CCl_4_+NAHP (Blue) (C). Data at each time point were from 9 mice. *ANOVA test P<0.05 compared to time 0 (before injection). (D) The means±SD of circulating histones at each week after CCl_4_ injection during a 28-day period (9 mice per group per time point). *ANOVA test, P<0.05 comparing to time 0 (before first injection). ^#^P=0.02 when compared the two groups at this time point. The means±SE of blood ALT levels (E) and liver injury scores (F) from 9 mice each group after 4 week treatments are presented. *ANOVA test P<0.05 compared to mice without treatment (Before). ^#^P<0.05 when compared to CCl_4_ group.

### Extracellular histones stimulate HSCs to increase the production of collagen I

Extracellular histones are recognized ligands for TLR receptors, including TLR2 and TLR4 ^[40, 45]^, both of which signal via the downstream MyD88-NFkB pathway. During liver fibrogenesis, HSCs express TLR4 receptor and its signalling enhances the activation of fibroblasts^[17]^. We directly treated LX2, a human HSC cell line with calf thymus histones. We found that histone concentrations between 2-10 μg/ml could significantly increase collagen I production. Five μg/ml histones incubated for 6 days were the optimal condition. This increased collagen I release into the culture media by nearly 2.5 fold and increased this value by 2.7 fold in cell lysates (Figure 2A and 2B). Concentrations of 20 μg/ml histones caused cell death in this culture condition (Data not shown). Using immunofluorescent staining, we found that LX2 cells treated with 5μg/ml histones were positively stained with anti-collagen I or anti-α-SMA compared to LX2 cells treated with no histones (Figure 2C), supporting that LX2 cells were activated by histone treatment and started to produce collagens.

**Figure 2.**
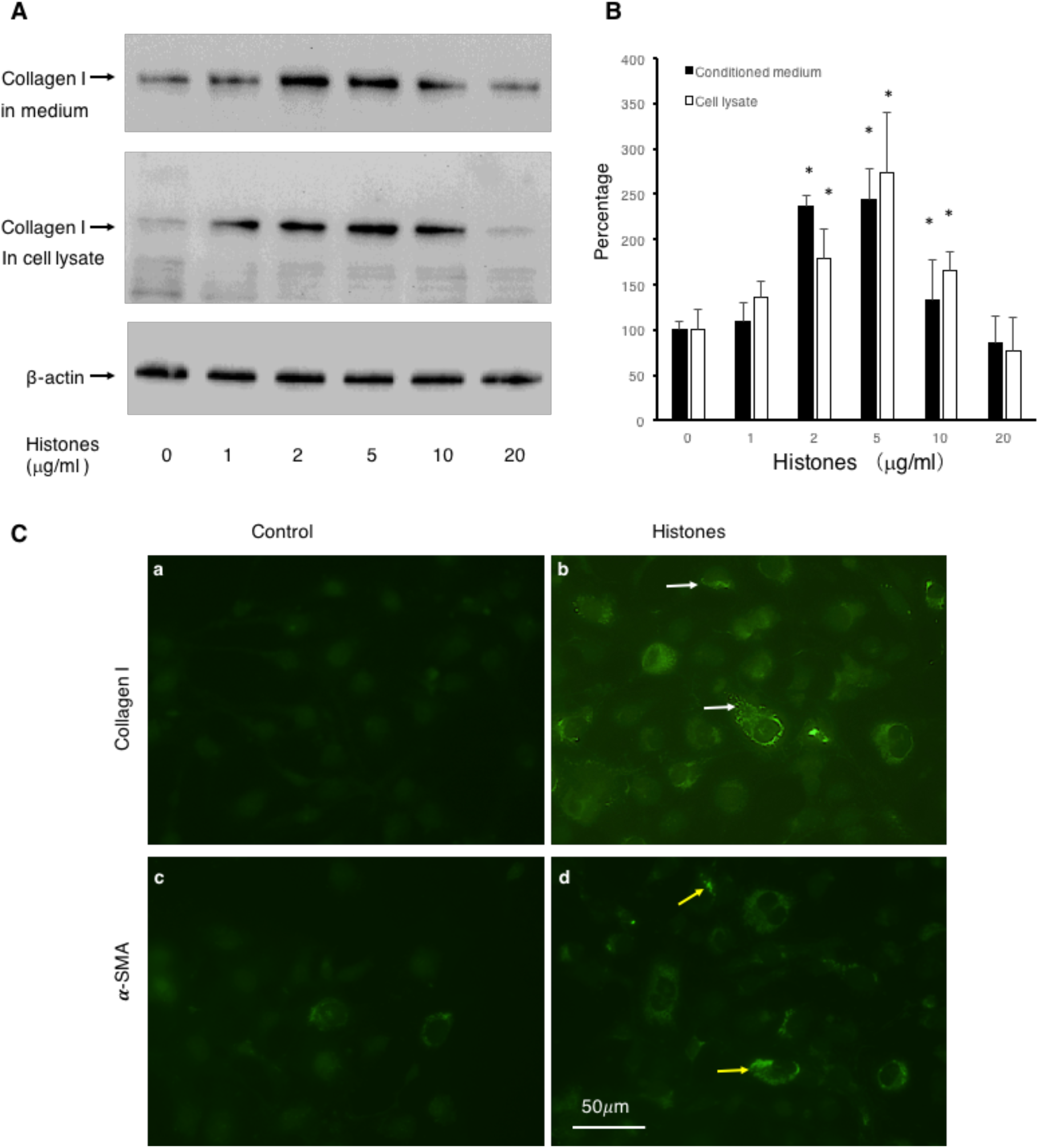
Extracellular histones induced collagen I production in LX2 cells. (A) Typical Western blots of collagen I in medium (upper panel) and lysates (middle panel) of LX2 cells treated with different concentrations of histones at day 6. Lower panel: beta actin in the cell lysates. (B) The means ±SD of the relative percentages of collagen/actin ratios with untreated (0 μg/ml histones) LX2 set at 100% from 5 independent experiments. ANOVA test, *P<0.05 compared to untreated. (C) Immunofluorescent staining of LX2 cells with anti-collagen I and anti-α-SMA antibodies. Control: cells were treated with routine medium without histones for 6 days. Histones: cells were treated with routine medium + histones 5μg/ml. Typical images are shown. White arrows indicate staining for collagen I, yellow arrows indicate staining for α-SMA. Bar=50μm.

### NAHP reduces histone-enhanced collagen production in LX2 cells and liver fibrosis induced by CCl_4_

Non-coagulant heparin (NAHP) has been demonstrated to bind histones and block their cytotoxicity^[26, 32, 49]^. In this study, we added NAHP (25 and 50 μg/ml) to culture medium to neutralize 5μg/ml calf thymus histones (approximately 1:2, and 1:4 molar ratio) and found that NAHP inhibited histone-enhanced collagen I synthesis in LX2 cells (Figure 3A, 3B). In H&E stained liver sections from the CCl_4-_induced mouse liver fibrosis model, CCl_4_ caused extensive hepatocytes damage compared to controls (Figure 3C, panel a), including hepatocyte swelling, vacuole formation, necrosis, Kupffer cell proliferation and immune cell infiltration (Figure 3C, panel b). NAHP intervention significantly reduced CCl_4_-induced hepatocyte vacuolization, necrosis and immune cell infiltration (Figure 3C, panel c). Immunohistochemical staining (IHC) with anti-α-SMA showed a larger area of positive staining in CCl_4_ treated mice compared to controls (Figure 3C, panel d and e). In NAHP treated mice, the CCl_4_-induced α-SMA staining was reduced (Figure 3C, panel f). Using Sirius red staining to visualise collagen, only the vascular walls were stained positively in controls (Figure 3C, panel g). In contrast, large areas of positive staining appeared in CCl_4_-treated mice (Figure 3C, panel h) and NAHP intervention reduced the areas of positive staining (Figure 3C, panel I). Using Image J software to quantify the areas of positive staining for α-SMA (Figure 3D) and for collagen (Figure 3E), we demonstrated that NAHP significantly reduced both α-SMA and collagen production. Using a colorimetric assay, hydroxyproline in liver tissues was detected and NAHP was shown to reduce CCl_4_-induced hydroxyproline elevation significantly (Figure 3F).

**Figure 3.**
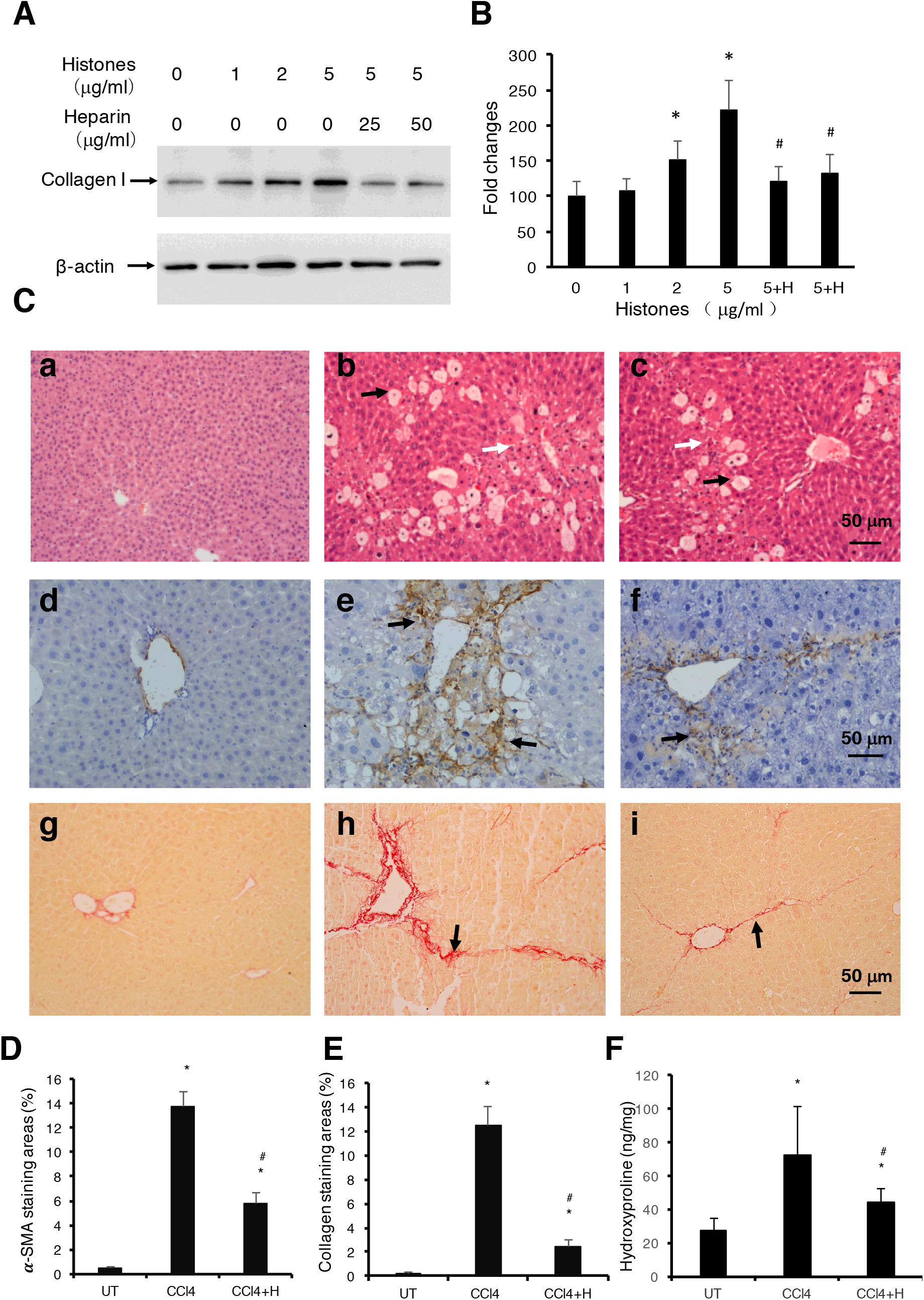
Effect of anti-histone reagent, NAHP, on histone-enhanced collagen I production in LX2 cells and CCl_4_-induced fibrosis in mice. (A) Typical Western blots of collagen I in LX2 cells treated with histones or histones+NAHP. Beta-actin is shown as loading reference. (B) The means ±SD of the relative percentage of collagen/actin ratios with untreated (0 μg/ml histones) LX-2 set at 100% from 3 independent experiments. ANOVA test, *P<0.05 compared to untreated. ^#^P<0.05 compared to 5μg/ml histones. (C)Typical images of stained liver sections (H&E staining: a, b, c; IHC staining with anti-α-SMA: d, e, f; and Sirius red staining: g, h,i) from normal mice (a, d, g); mice treated with CCl_4_ (b, e, h) and mice treated with CCl_4_+NAHP (CCl_4_+H) (c, f, i). Arrows in b and c: black indicate hepatocyte swelling and white indicate necrosis and immune cell infiltration. Arrows in e and f: indicate staining for smooth muscle actin. Arrows in h and i: indicate collagen deposition. (D) and (E) Means ±SD of percentage of areas of staining for α-SMA(D) and Sirius red staining for collagen (E) (9 mice per group, and 6 randomly selected sections per mice). ANOVA test *P<0.05 compared to control. ^#^P<0.05 when compared to CCl_4_ alone. (F) Means ±SD of hydroxyproline levels in liver tissues from 9 mice per group. ANOVA test *P<0.05 compared to control. ^#^P<0.05 when compared to CCl_4_ alone.

### TLR4 is involved in histone-enhanced collagen production in LX2 cells and in liver fibrosis induced by CCl_4_

To understand how histones are involved in liver fibrosis, we specifically tested if histones when activating the TLR/MyD88 signalling pathway, played a major role in this process; this was based on previous reports that TLR4 and MyD88 were involved in fibrogenesis^[17, 19]^. We used human TLR4-neutralizing antibodies (TLR4i) to treat LX2 cells and found that histone-enhanced collagen I production was significantly reduced (Figure 4A). To confirm that the TLR4-MyD88 signalling pathway is actually involved in CCl_4_-induced liver fibrosis, TLR4 and MyD88 gene knockout mice were used. We found that the extent of fibrosis was significantly less in both TLR-4 (-/-) and MyD88(-/-) mice than in wt mice treated with CCl_4_ (Figure 4B and 4C). These results are consistent with the previous publication that demonstrated TLR-4 and MyD88 deficiency reduced and liver fibrosis in the mouse bile duct ligation model^[50]^. These findings strongly suggest that extracellular histones may activate the TLR4-MyD88 signalling pathway to enhance collagen I production and liver fibrosis. However, further investigation is required to substantiate this point.

**Figure 4.**
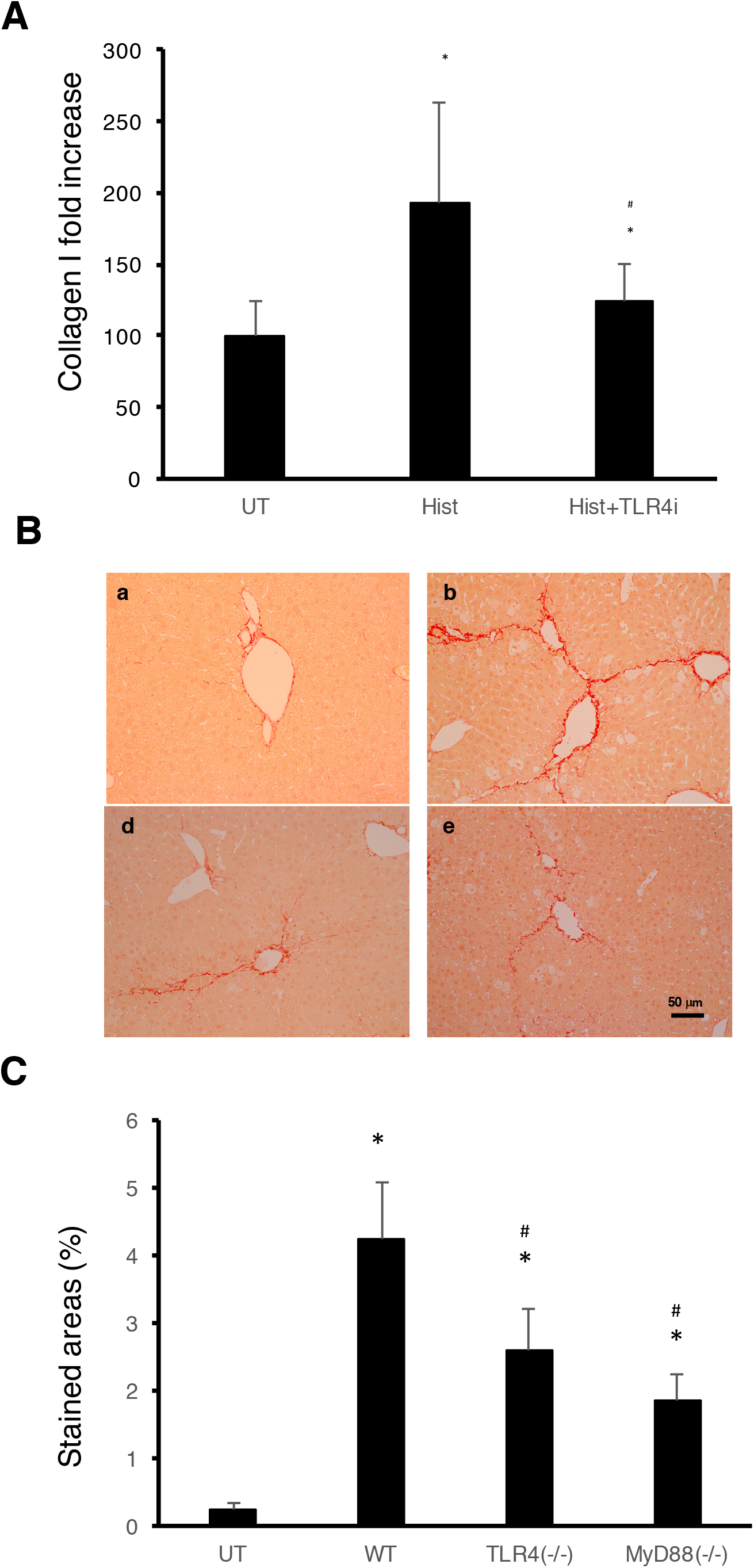
TLR4 is involved in histone-enhanced collagen I production and CCl_4_-induced mouse liver fibrosis. (A) LX2 cells were treated with 5μg/ml histones (Hist) in the presence or absence of TLR4 neutralizing antibody (TLR4i). The means±SD of the relative percentage of collagen I/beta-actin ratios are presented with control (UT) set at 100% from 3 independent experiments. ANOVA test *P<0.05 compared to UT. ^#^P<0.05 when compared to histone alone. **(B)** Typical images of Sirius red staining of liver section from untreated wt C57BL/j mice (a), CCl_4_-treated wt mice (b), TLR-2 (c), TLR-4 (d) and MyD88 (e) knockout mice are presented. (C) Means±SD of stained areas of liver sections from untreated wt mice (UT), CCl_4_-treated wt mice (WT), CCl_4_-treated TLR4(-/-) (TLR4 (-/-)) and MyD88(-/-) (MyD88(-/-)) knockout mice. Eight mice were in each group and 6 slides from each mouse were analysed. ANOVA test *P<0.05 compared to untreated wt mice (UT). ^#^P<0.05 compared to CCl_4_-treated wt mice.

## DISCUSSION

This study has proposed a novel concept that extracellular histones act as a pathogenic factor in liver fibrosis. High levels of circulating histones in CCl_4_-treated mice have been detected from 4 hours after first administration of CCl_4_ and remain high during the duration of fibrosis induction. *In vitro*, extracellular histones directly enhanced collagen I production in LX2, a human HSC, by up to 2.7 fold. When NAHP was used to neutralize histones, it significantly reduced histone-enhanced collagen I production in LX2 cells and significantly reduced fibrosis in CCl_4_-treated mice. When TLR4 neutralizing antibody was used, histone-enhanced collagen I production in LX2 cells was also reduced. Moreover, when CCl_4 was_ injected into TLR4 or MyD88 knockout mice, they showed significantly less fibrosis compared to wt mice, suggesting that both TLR4 and MyD88 were involved in histone-enhanced fibrosis.

TLRs not only recognize pathogen-associated molecular patterns (PAMPs) from various microbial infections, but also respond to DAMPs from host cell damage^[51]^. TLR4 recognizes LPS from Gram negative bacteria as well as histones released after cell death. TLR4 can activate NFkB via MyD88^[40, 44, 52, 53]^. The TLR4/MyD88 pathway has been demonstrated to promote liver fibrosis in a mouse bile duct ligation model by enhancing TGF-β signalling^[50]^. LPS as a TLR-4 ligand was elevated 3-6 fold in this model probably due to obstruction of bile ducts and subsequently bacterial infection^[50]^. However, circulating LPS concentration did not increase in the CCl_4_ mouse model ^[54]^. Without infection, LPS from digestive tracts could enter the circulation in the late stage of fibrosis or cirrhosis, whilst the wound-healing process is initiated immediately after injury. Therefore, LPS is unlikely to be an initiating factor in fibrosis. Extracellular histones are the newly-found TLR4 ligands and are most likely to mediate the initial stage of fibrogenesis by activating the TLR4-MyD88 signalling pathway.

The levels of extracellular histones required for stimulating LX2 human HSCs cells are between 2-10 μg/ml, which is much lower than that detected in the circulation of mice, [median, quartiles]: 70.5μg/ml [19.5, 246.9] between 8-48h after CCl_4_ administration. This is due to the severe and extensive damage to hepatocytes caused by the radical CCl_3_, a metabolite of CCl_4_. The highest levels of histones are comparable to what we detected in critical illnesses^[26, 31, 32]^ and are most likely able to cause further liver injury^[55]^. However, plasma can neutralize histones at a concentration of up to 50μg/ml *in vivo* or *ex vivo* making cells more resistant than those in cultured medium^[32]^. In addition, in our mouse model, the interval between 2 doses of CCl_4_ was 84h, therefore more than half of this period had histone levels around 50μg/ml, which may be sufficient in maintaining TLR4 activation. Elevation of circulating histones was also found in many types of liver disease and the high levels of circulating histones could cause secondary liver injury^[26, 40, 56]^. Some of human chronic liver diseases, such as chronic hepatitis B and C, alcoholic and autoimmune liver diseases may have low but constant circulating histones to serve as alarmins, which signal tissue injury and initiate repair. However, this needs further clinical investigation.

The NAHP could bind to histones, but will not lower the levels of circulating histones. In the first week, NAHP seems to increase the levels of circulating histones in the CCl_4_-treated mice and this could be explained by its ability to reduce histone clearance as we have observed previously^[26]^. NAHP binding to extracellular histones prevents their binding to the cell plasma membrane and other targets, such as prothrombin^[32–34]^ and thereby detoxifies histones. In this study, we found that ALT and liver injury scores were significantly reduced by NAHP in CCl_4_-treated mice, strongly indicating that high levels of circulating histones are toxic to liver cells and their toxicity can be neutralized by NAHP. This protective effect may also contribute to the reduction of liver fibrosis.

Heparin was reported to reduce collagen fibre formation and liver fibrosis two decades ago^[57–59]^. Enzymatically depolymerized Low Molecular Weight Heparins were shown to inhibit CCl_4_-induced liver fibrosis by reducing TNFα and interleukin (IL)-1β^[60]^. Although extracellular histones could enhance cytokine release, including IL-6, IL-1β, and TNFα^[33, 40]^, we suggest that the major mechanism for heparin-inhibited liver fibrosis is most likely due to its blocking of extracellular histones’ ability to activate the TLR4-MyD88 signalling pathway. Heparin has been used as an anticoagulant for many decades and overdoses may cause bleeding, which can be avoided by using NAHP^[26]^. In addition, the NAHP alone appears to have no significant toxicity for liver, but it significantly reduced histone-enhanced collagen I expression in LX2 cells and reduced CCl_4_-induced liver fibrosis. Therefore, this novel finding may help advance translating this old discovery into a clinical application.

There are a few limitations in this study. Firstly, we only demonstrated that extracellular histones could directly stimulate HSC to produce collagen I, which could be reduced by NAHP or TLR4 neutralizing antibody, and proposed a novel mechanism of enhanced collagen production by HSC. However, only one type of animal model for liver fibrosis was used and this may not truly represent the fibrosis caused by different diseases. However, it is sufficient to demonstrate a novel concept. Secondly, liver fibrosis is a complicated process involving many types of cells and signalling pathways. In our in vivo study, no direct link has been demonstrated between histones and binding to TLR4. In addition, the NAHP may have other effects besides blocking histones to activate TLR4 and high levels of circulating histones may also affect many other types of cell besides HSC. Therefore, further mechanistic studies in vivo are required and development of better reagents to target histones and potential signalling pathways may offer more effective therapies in the future.

## Author Contribution

Wang Z conceived the study; Cheng ZX assisted animal experiments and hydroxyproline measurement; Lin Z and Abrams ST assisted in performing in vitro experiments; Yates ED synthesized and characterised non-anticoagulant heparin. Abrams helped edit figures. Yu Q, Yu WP, Chen PS, Toh CH and Wang GZ supervised the work and were involved in data analysis and manuscript writing. All authors have read and agreed to the published version of the manuscript.

## Acknowledgement

Thanks to all the staff in animal house for their support and to the technicians in the pathology laboratory for technical support and clinical biochemistry for ALT measurement in Zhongda Hospital, Nanjing, China. Thanks to Professor Philip Rudland in the University of Liverpool for his critical reading and correction of the manuscript. Thanks to Dr. Steven Lane in the Department of Statistics, University of Liverpool for his assistance in statistical analysis.

